# Elucidating Fibroblast Growth Factor-induced kinome dynamics using targeted mass spectrometry and dynamic modeling

**DOI:** 10.1101/2023.01.27.525819

**Authors:** Tim S. Veth, Chiara Francavilla, Albert J.R. Heck, Maarten Altelaar

**Affiliations:** Biomolecular Mass Spectrometry and Proteomics, Bijvoet Center for Biomolecular Research and Utrecht Institute for Pharmaceutical Sciences, University of Utrecht, Padualaan 8, Utrecht, 3584 CH, The Netherlands; Netherlands Proteomics Center, Padualaan 8, Utrecht, 3584 CH, The Netherlands; Division of Molecular and Cellular Function, School of Biological Science, and Manchester Breast Centre, Manchester Cancer Research Centre, Faculty of Biology Medicine and Health (FBMH), The University of Manchester, M139PT, Manchester, UK

**Keywords:** Phosphoproteomics, kinase, targeted mass spectrometry, kinase dynamics, fibroblast growth factors, dynamic modeling, fibroblast growth factor receptor, breast cancer

## Abstract

Fibroblast growth factors (FGFs) are paracrine or endocrine signaling proteins that, activated by their ligands, elicit a wide range of health and disease-related processes, such as cell proliferation and the epithelial-to-mesenchymal transition (EMT). The detailed molecular pathway dynamics that coordinate these responses have remained to be determined. To elucidate these, we stimulated MCF-7 breast cancer cells with either FGF2, FGF3, FGF4, FGF10, or FGF19. Following activation of the receptor, we quantified the kinase activity dynamics of 44 kinases using a targeted mass spectrometry assay. Our system-wide kinase activity data, supplemented with (phospho)proteomics data, reveal ligand-dependent distinct pathway dynamics, elucidate the involvement of not earlier reported kinases such as MARK, and revise some of the pathway effects on biological outcomes. In addition, logic-based dynamic modeling of the kinome dynamics further verifies the biological goodness-of-fit of the predicted models and reveals tight regulation of the RAF kinase family.

## Introduction

Fibroblast growth factors (FGFs) and co-factors heparin/heparin-sulfate or beta-klotho induce trans-autophosphorylation upon binding to fibroblast growth factor receptors (FGFRs), thereby activating signaling pathways and regulating diverse biological processes (Ornitz & Itoh, 2022; Sarabipour & Hristova, 2016; Su et al., 2014) (Kuro-o, 2019; Spivak-Kroizman et al., 1994). There are 18 FGF ligands known so far that can activate the 7 alternatively spliced isoforms of 4 FGFR genes. Specific combinations of receptor and ligand result in the regulation of a plethora of diverse cellular processes, including cell differentiation, cell proliferation, and epithelial-mesenchymal transition (EMT) (Chen, 2005; Xie et al., 2020).

Besides the role of FGFs in health, during development and adult life, dysregulated FGF-FGFR signaling is implicated in various types of cancer, including breast cancer (Francavilla & O’Brien, 2022; Korc & Friesel, 2009; Presta et al., 2017; Y. Zhou et al., 2020). FGF2 is commonly detected in the tumor microenvironment of breast cancer and can induce tumor growth (Giulianelli et al., 2019; Sharpe et al., 2011). FGF3, FGF4, and FGF19 are located on the 11q13 amplicon, which is amplified in 15-20% of breast cancer patients and are all linked to increased tumor progression (Karlsson et al., 2011, p. 201; W. Wang et al., 2015; C. Zhang et al., 2020; Zhao et al., 2018). FGF10 can drive type III EMT in breast cancer, promoting invasiveness (Abolhassani et al., 2014). These unfavorable effects in breast cancer patients result from diverse and complex FGF-driven cellular signaling (Presta et al., 2017).

The fine-tuned coordination of the diverse FGF-driven cellular processes is thought to be regulated by the MAPK/ERK pathway, the PI3K pathway, the PLCγ pathway, and the JAK-STAT pathway (Dailey et al., 2005; Ferguson et al., 2021) (Ornitz et al., 1996; Touat et al., 2015; X. Zhang et al., 2006). For example, the MAPK/ERK pathway is thought to drive cell proliferation, and the PI3K pathway is believed to regulate EMT (Katoh & Nakagama, 2014; Tomita et al., 2021). These pathways are highly dependent on multiple kinases that relay signals by adding phosphate groups to proteins or other molecules. Kinase activity is often determined by phosphorylation in the kinase activation loop, which can be measured and quantified using a targeted mass spectrometry based kinome assay (Nolen et al., 2004; Schmidlin et al., 2019). Even though the main pathways involved in FGF signaling are elucidated, molecular mechanistic insights into the regulations of the differential cellular processes are still largely lacking (Gurzu et al., 2019; Ramos et al., 2010).

FGF2, FGF3, FGF4, FGF10, and FGF19 are all associated with breast cancer, however, insights into the differential signaling of these FGFs are lacking. It is unclear what pathways and kinases are regulated by the different FGFs. Also, no mechanistic signaling comparisons are investigated to elucidate the importance of each of the FGFs and their possible roles in breast cancer. Gaining these biological insights is key to understanding the implications of FGF signaling in breast cancer.

Here, we aim to broaden our understanding of FGF signaling by quantifying temporal kinase activation dynamics using a selected reaction monitoring assay (SRM) with broad coverage of kinases that are involved in the FGFR signaling pathway. To verify the biological results from the longitudinal SRM data, we created a dynamic mechanistic model of the signaling pathway using logic-based ordinary differential equations. To explain discrepancies in our developed model, we used modeling-guided analysis of shotgun phosphoproteomics data. Our approach successfully mapped FGF2, FGF3, FGF4, FGF10, and FGF19 signaling in breast cancer cell lines and allowed us to add hitherto unknown involved kinases and signaling dynamics to FGF stimulations.

## Methods

### Cell culture

MCF-7 (ATCC), BT-474 (ATCC), and EFM-192a (DSMZ) cells were grown in Dulbecco’s modified Eagle’s medium (DMEM) supplemented with 10% FBS (Sigma) and 2mM glutamine. Cells were regularly tested for mycoplasma. All cells were cultured in a humidified incubator equilibrated with 5% CO2 at 37 °C. Experiments were performed after the 5^th^ passage and before the 20^th^ passage to limit cell heterogeneity between experiments.

### Sample preparation for mass spectrometry

For mass spectrometry experiments, ∼5 million cells were plated in triplicates in 10cm plates in regular medium. After 24 hours, the medium was changed to serum-starved medium supplemented with 5 µg/mL heparin (Thermo Scientific). After 24 hours, cells were incubated with 50 ng/mL of either FGF2 (Peprotech), FGF3 (KyvoBio), FGF4 (Peprotech), FGF10 (Peprotech), or FGF19 (Peprotech). Cells were washed three times with ice-cold PBS, scraped, and snap-frozen until further sample preparation.

### Cell growth assay

Triplicate groups of ∼0.1 million cells were plated in 12 well-plates again first in regular medium and subsequently in medium with either 5 µg/mL heparin or without. After 24 hours, one of the 5 different FGF ligands was added, and the plate was incubated in an IncuCyte ZOOM™ at 37 °C/5% CO2 until the end of the experiment. Pictures of each well were taken every hour, of which the percentage plate coverage was determined. Significance between groups was determined using an ANOVA and Tukey’s range test (p < 0.05).

### Scratch wound healing assay

In 12 well plates, triplicates of 3e5 cells were plated in a regular medium, after 24 hours, the medium was changed to starved medium supplemented with 5 µg/mL heparin. Subsequently, the cells were verified to be confluent when the scratch assay was performed (Liang et al., 2007). The scratch assay was analyzed as described before (Suarez-Arnedo et al., 2020). In short, using the ImageJ/Fiji script “Wound Healing Size Tool”, the percentage of wound closure was calculated between t = 24h and t = 0 (Schindelin et al., 2012). Significance between groups was determined using an ANOVA and Tukey’s range test (biological triplicates, p < 0.05).

### Spectral library generation

Spectral libraries were used to determine peptide fragmentation characteristics and their indexed retention time, which are key for identifying peptides in the tier 2 SRM assay. The custom mix of heavy labeled peptides (JPT or ThermoFisher Scientific) was mixed with iRT peptides (Biognosys) and analyzed using an Orbitrap Q-Exactive HF (ThermoFisher Scientific). An unscheduled parallel reaction monitoring (PRM) method scanned for the +2 and +3 charged peptides, including all possible methionine oxidations. Peptides were separated using a 2 h gradient a 120k resolution was used for the PRM assay, resulting in a minimum of 5 spectra per peptide. Raw files were analyzed using MaxQuant (version 1.6.10.43), carbamidomethyl cysteine as fixed modification, and the variable modifications serine/threonine/tyrosine phosphorylation, methionine oxidation, and isotope labels. The search results were filtered using a 1% FDR cut-off, and subsequently, using Skyline (version 20.1.1.83), pseudo-MS2 spectra were generated, which were used as the peptide library.

### SRM assay development

The SRM assay was developed using previously described methods (Schmidlin et al., 2019). The assay was developed on a TSQ Altis (ThermoFisher Scientific). In brief, the 10 most intense fragment ions from the library were used as initial transitions. These transitions were used to optimize multiple parameters, such as retention time and collision energy. The collision energy was optimized per transition using Skyline, with the TSQ Vantage CE formula as starting point (CE = 0.03 m/z + 2.905 for doubly charged precursors and CE = 0.038 m/z + 2.281 for precursor charges of three and higher) and optimized using steps of 1 voltage.

### Protein digestion selected reaction monitoring assay

Snap-frozen protein pellets were lysed, reduced, and alkylated in lysis buffer (1% sodium deoxycholate (SDC), 10 mM tris(2-carboxyethyl)phosphine hydrochloride (TCEP)), 40 mM chloroacetamide (CAA), and 100 mM TRIS, pH 8.0 supplemented with phosphatase inhibitor (PhosSTOP, Roche) and protease inhibitor (cOmplete mini EDTA-free, Roche). Cells were heated at 95C and sonicated with a Bioruptor Plus (Diagenode) for 15 cycles of 30 s. Bradford protein assay (Bio-Rad Protein Assay Kit I, Bio-Rad) was used to determine the protein amount, after which samples were split into 200µg aliquots. Proteins were digested overnight at 37C with trypsin (1:50 µg/µg) (Sigma-Aldrich) and lysyl endopeptidase (1:75 µg/µg) (Wako). Heavy labeled phosphopeptides were added to the samples. The SDC was precipitated with 2% formic acid (FA) twice, after which samples were desalted and enriched in an automated fashion using the AssayMap Bravo platform (Agilent Technologies) with corresponding AssayMap C18 (Agilent Technologies) reverse-phase column as previously described (Post et al., 2017).

### SRM LC-MS/MS Setup

Samples were analyzed on a TSQ Altis (Thermo Scientific) coupled to an UltiMate 3000 (Thermo Scientific), and an easy spray analytical column (ES802A, 25 cm, 75 mm ID PepMap RLSC, C18, 100 A°, 2 mm particle size column (Thermo Scientific)). First, samples were reconstituted in 2% LC-MS grade formic acid. Samples were loaded on a trap column (Acclaim™ PepMap™ 100 C18 HPLC Column 0.3×5mm with 5 μm particles (Thermo Scientific)) with 2.2% Buffer A (0.1% FA) for 3 minutes and subsequently separated using 0-32% buffer B (99.9%ACN, 0.1%FA) in 35 min at 300nL/min and followed by a 20 min column wash with 80% buffer B at 300nL/min, and 10-minute column equilibration at 2.2% B. The TSQ Altis spray voltage was set at 1.9 kV and fragmented at 1.5 mTorr in the second quadrupole. The first quadrupole was set at 0.7 da FWHM, and the third quadrupole at 1.2 da FWHM. All transitions were measured with optimized collision energy without scheduling and a cycle time of 1.5 sec.

### SRM data assessment

All experiments were analyzed using Skyline-Daily (version 20.2.1.404) (Pino et al., 2020). The quality of the peptides was assessed mainly on the signal similarity between the heavy and the light peptides. The most important aspects were perfect co-elution, peak shape, and relative contributions of each transition between the heavy and the light peptide. A rdotp > 0.95 was maintained to indicate the similarity between the heavy and the light peptide. In-house R scripts were used for further data visualization and analysis.

### Logic-based dynamic modeling

Logic-based dynamic modeling was performed as described earlier (Tognetti et al., 2021). In short, first, a prior knowledge network (PKN) was generated using Omnipath and converted to a simple interaction file (SIF) (Türei et al., 2016). Normalization was done per kinase across all the FGFs. The average fold change to t=0 was scaled between 0-1 using the 99% interquartile range (biological triplicates) described in **Equation 1**.

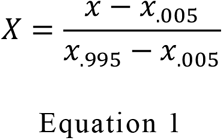

Values < 0 or > 1 were set to 0 or 1, respectively. The different FGFs were set to 0.75 for their modeling.

The model was trained using the freely available CNORode for all FGFs simultaneously (Terfve et al., 2012). Each kinase can be described using a continuous update function B_i_ where the activity of a kinase x_i_ is predicted {0,1} using the associated upstream effectors, as shown in Equation 2 (Wittmann et al., 2009).

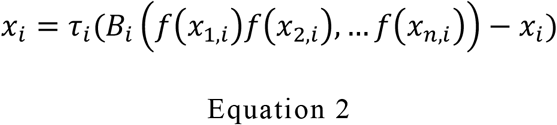

*τ*_*i*_ can be interpreted as the kinase responsiveness to upstream effectors where a small value indicates a slower response. Each transfer function is a Hill-type function, as previously described and presented in Equation 3 (Eduati et al., 2017).

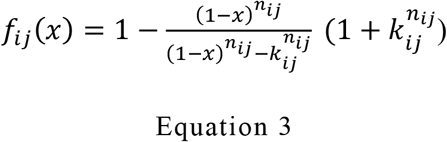

The sigmoidal shape curve is determined by parameters n and k. The k parameter can be interpreted as the strength of the interaction where a high k value describes a high signal throughput.

### Kinase dynamic parameter estimation

Each kinase is assigned a fixed n value of 3 and a k and τ value determined by the dynamic modeling. CNORode and the MEIGOR toolkit were used, which uses the normalized kinase activity data and the PKN to determine the best k and τ values based on the smallest root-mean-square error (RMSE) (Egea et al., 2014). The method entails L2 normalization to prevent overfitting, which was set to a value of 10^−5^. The update function was verified to have achieved optimal performance based on the RMSE response curves. Model goodness of fit was determined using Pearson’s r and the RMSE of all measured and predicted time points of all kinases. The biological RMSE was determined using the deviation between the measured values and the mean.

### Peptide work-up untargeted phosphoproteomics

Peptide work-up was performed identically to the SRM peptide workup except that no heavy labeled peptides were added after digestion.

### Peptide work-up untargeted proteomics

Snap-frozen protein pellets were lysed, reduced, and alkylated in lysis buffer (1% sodium deoxycholate (SDC), 10 mM tris(2-carboxyethyl)phosphine hydrochloride (TCEP)), 40 mM chloroacetamide (CAA), and 100 mM TRIS, pH 8.0 supplemented with protease inhibitor (cOmplete mini EDTA-free, Roche). Cells were heated at 95C and sonicated with a Bioruptor Plus (Diagenode) for 15 cycles of 30 s. Bradford protein assay (Bio-Rad Protein Assay Kit I, Bio-Rad) was used to determine the protein amount, after which samples were split into 10µg aliquots. Proteins were digested overnight at 37C with 1:50 trypsin (Sigma-Aldrich) and 1:75 and lysyl endopeptidase (Wako), after which samples were desalted using an Oasis® platform, dried down, and stored at -80 until further use.

### Data-dependent analysis of untargeted phosphoproteomics

Samples were suspended in 2% formic acid and analyzed on an Exploris (Thermo Scientific) coupled to an UltiMate 3000 (Thermo Scientific), fitted with a µ-precolumn (C18 PepMap100, 5µm, 100 Å, 5mm × 300µm; Thermo Scientific), and an analytical column (120 EC-C18, 2.7µm, 50cm × 75µm; Agilent Poroshell). Peptides are loaded in 9% Buffer A (0.1% FA) for 1 minute and separated using 9-36% buffer B (80%ACN, 0.1%FA) in 97 min at 300nL/min and followed by a 6 min column wash with 99% buffer B at 300nL/min, and a 10-minute column equilibration at 9% B. The MS was operated in DDA mode, with the MS1 scans in a range of 375-1600 m/z acquired at 60k, using an automatically set AGC target. MS2 scans were acquired with a 16s dynamic exclusion at a 30k resolution, 28% normalized collision energy, and an isolation window of 1.4 m/z.

Raw files were processed via MaxQuant version 1.6.17.0 using the verified human proteome from UniprotKB (release 09-2019) containing 20369 proteins (Tyanova et al., 2016). A maximum of 5 modifications and two miscleavages were set using fixed carbamidomethyl modification, and the variable modifications oxidized methionine, protein N-terminal acetylation, and serine/threonine/tyrosine phosphorylation. The protein and peptide false discovery rates were set to < 0.01 and conducted with match between runs enabled. No normalization or imputation was applied.

### Shotgun proteomics analysis

Samples were suspended in 2% formic acid and analyzed on a Q-Exactive HF (Thermo Scientific) coupled to an UltiMate 3000 (Thermo Scientific), fitted with a µ-precolumn (C18 PepMap100, 5µm, 100 Å, 5mm × 300µm; Thermo Scientific), and an analytical column (120 EC-C18, 2.7µm, 50cm × 75µm; Agilent Poroshell). Peptides are loaded in 9% Buffer A (0.1% FA) for 1 minute and separated using 9-44% buffer B (80%ACN, 0.1%FA) in 155 min at 300nL/min and followed by a 6 min column wash with 95% buffer B at 300nL/min, and a 10-minute column equilibration at 9% B. The MS was operated in DDA mode, with the MS1 scans in a range of 375-1600 m/z acquired at 60k, using an AGC target of 3e6. MS2 scans were acquired with a 24s dynamic exclusion at a 30k resolution, 27% normalized collision energy, and an isolation window of 1.4 m/z.

Raw files were processed via MaxQuant version 1.6.17.0 using the verified human proteome from UniprotKB (release 09-2019) containing 20369 proteins (Tyanova et al., 2016). A maximum of 5 modifications and 2 miscleavages was set using fixed carbamidomethyl modification, and the variable modifications oxidized methionine and protein N-terminal acetylation. The protein and peptide false discovery rates were set to < 0.01 and conducted with match between runs enabled. Further analysis was performed using artMS version 1.12.0 building on MSstats (Choi et al., 2014; Jimenez-Morales et al., 2019). MSstats imputation was done using accelerated failure time modeling, and the samples were median normalized after imputation.

### FGFR qPCR quantification

MCF-7 cells were plated in triplicates. Subsequently, the samples were lysed and prepared for qPCR analysis using the protocol adapted from (Korsten et al., 2022). In short, samples were lysed and isolated using the vendor’s instructions of NucleoSpin RNA plus (Macherey-Nagel) with the addition of a DNase removal step using RNase-Free Dnase (Qiagen). Next, 500 ng of RNA was used to obtain cDNA using the vendor’s instructions of AH iScript (Bio-Rad). The qPCR was performed at 95C for 10min, followed by (95C for 30 sec, 55C for 30 sec, and 72C for 30sec), a total of 40 times. Normalization was performed using Beta-actin and Glyceraldehyde-3-phosphate dehydrogenase following (Taylor et al., 2019). The primers used are listed in supplementary table 1.

### RAP1 activation assay

RAP1 activity was determined following the supplier’s instructions (Merck, Cat# 17-321). In short, MCF-7 cells were plated in 15cm plates and incubated for 60 minutes with FGF2, FGF3, FGF4, FGF10, FGF19, or without FGF and lysed using the provided lysis buffer. After, equal amounts of protein were used for the RAP1 pulldown, including one positive control consisting of MCF-7 cell lysates incubated with GTPγS. Subsequently, a western blot was conducted using the provided RAP1 antibodies. Linear adjustments were performed using Fiji (Schindelin et al., 2012).

## Results

### Dynamic kinase activity quantification

Here we performed (phospho)proteomics experiments to elucidate the specific effect of different FGF ligands on FGFR activation and downstream signaling. Thereby we focused on FGFR signaling in breast cancer cells induced by either FGF2, FGF3, FGF4, FGF10, or FGF19. To understand signaling, we quantified temporal system-wide kinase activity using a dedicated selected reaction monitoring (SRM) assay targeting the activation loops of a widespread panel of kinases (**Figure 1A**). To increase kinome coverage of the kinases involved in the FGFR signaling pathway, nearly 200 phosphopeptides spanning 50 kinases were included in the assay earlier developed by Schmidlin et al. (2019), resulting in an assay comprising 484 phosphopeptides on 197 kinases (**Table S1**) (Schmidlin et al., 2019).

**Figure 1.**
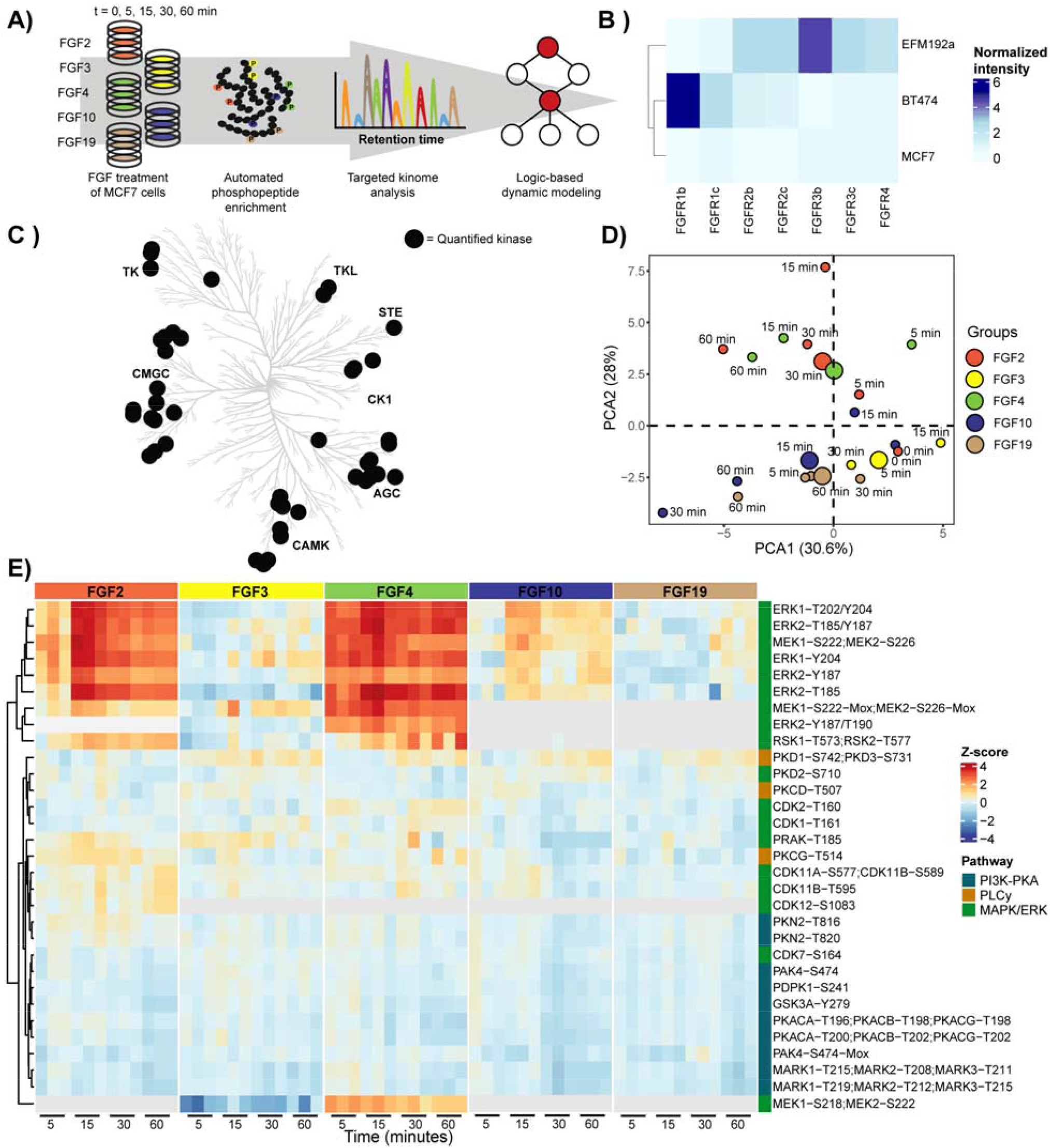
Stimulation with FGF2, FGF3, FGF4, FGF10, or FGF19 results in differential kinome regulation. **A)** Schematic overview of the experimental approach, whereby a targeted kinase activation loop SRM assay was used to monitor system-wide kinase activity upon treatment of MCF-7 cells with distinct FGF ligands. **B)** qPCR experiments were performed to monitor FGFR expression on three different cell lines. FGFR expression was normalized to Beta-actin and Glyceraldehyde-3-phosphate dehydrogenase. **C)** Kinome tree with kinases significantly regulated by at least one of the tested FGF ligands represented by black dots (ANOVA p < 0.05, triplicate measurements). Principle Component Analysis (PCA) of the kinase activity data at different time points and with the different tested FGFs. Mean values were used for the independent triplicate measurements. **E)** Heatmap through unsupervised hierarchical clustering of all significantly (ANOVA p < 0.05) regulated phosphorylated peptides over all time points and FGFs (with each experiment performed in triplicate). Not identified phosphorylated peptides are represented in grey.

To select an appropriate cell line, this SRM assay was initially used to monitor the system-wide kinome activity response of a set of breast cancer cell lines, namely MCF-7, BT-474, and EFM-192a cells, upon FGF2 and FGF4 stimulation as these bind the majority of FGFR spliceoforms. From these data, we concluded that MCF-7 cells especially displayed a broad kinome response after stimulation (**Figure S1**). We reasoned this would be explained by FGFR expression, however, surprisingly, qPCR quantification of FGFR expression in the panel of tested cells showed that the MCF-7 cells exhibited an overall low expression of FGFRs (**Figure 1B** and **Figure S2**). This highlights that FGFR expression alone does not solely determine the extent of downstream signaling. Due to the observed broad kinome response, we did proceed with the MCF-7 cells, which were incubated with either FGF2, FGF3, FGF4, FGF10, or FGF19, and the cofactor heparin for 0, 5, 15, 30, and 60 minutes (Eswarakumar et al., 2005; Wolf et al., 2008). Using the kinase activation loop SRM assays, we quantified kinase activity profiles of 46 phosphorylated sites spanning 44 kinases (**Table S2**). Of these, 35 kinases displayed significant regulation over time (ANOVA p < 0.05) upon stimulation with at least one of the 5 tested FGF ligands. Each of the tested ligands resulted in differential regulation of kinases across most kinase families (**Figure 1C**) that were, as expected, primarily members of the MAPK/ERK, PI3K, and/or PLCγ pathways (**Figure 1D/E**) (Ornitz & Itoh, 2015, 2022).

### Fine-tuned activation of the MAPK/ERK signaling pathway

As the MAPK/ERK pathway is known to be involved in FGF signaling, we first compared the kinase activity profiles acquired with the SRM assays of kinases involved in this pathway. FGF-stimulated MAPK/ERK activation is commonly regarded to be directed via the RAS-RAF-MEK-ERK signaling cascade (Azami et al., 2017; Bockorny et al., 2018; Cho et al., 2009; Kunath et al., 2007; Lovicu & McAvoy, 2001; Shalaby et al., 2009; Tomita et al., 2021). In MCF-7 cells, only FGF2, FGF4, and FGF10 treatments significantly activated several of the kinases in the MAPK/ERK pathway (**Figure 2A**).

**Figure 2.**
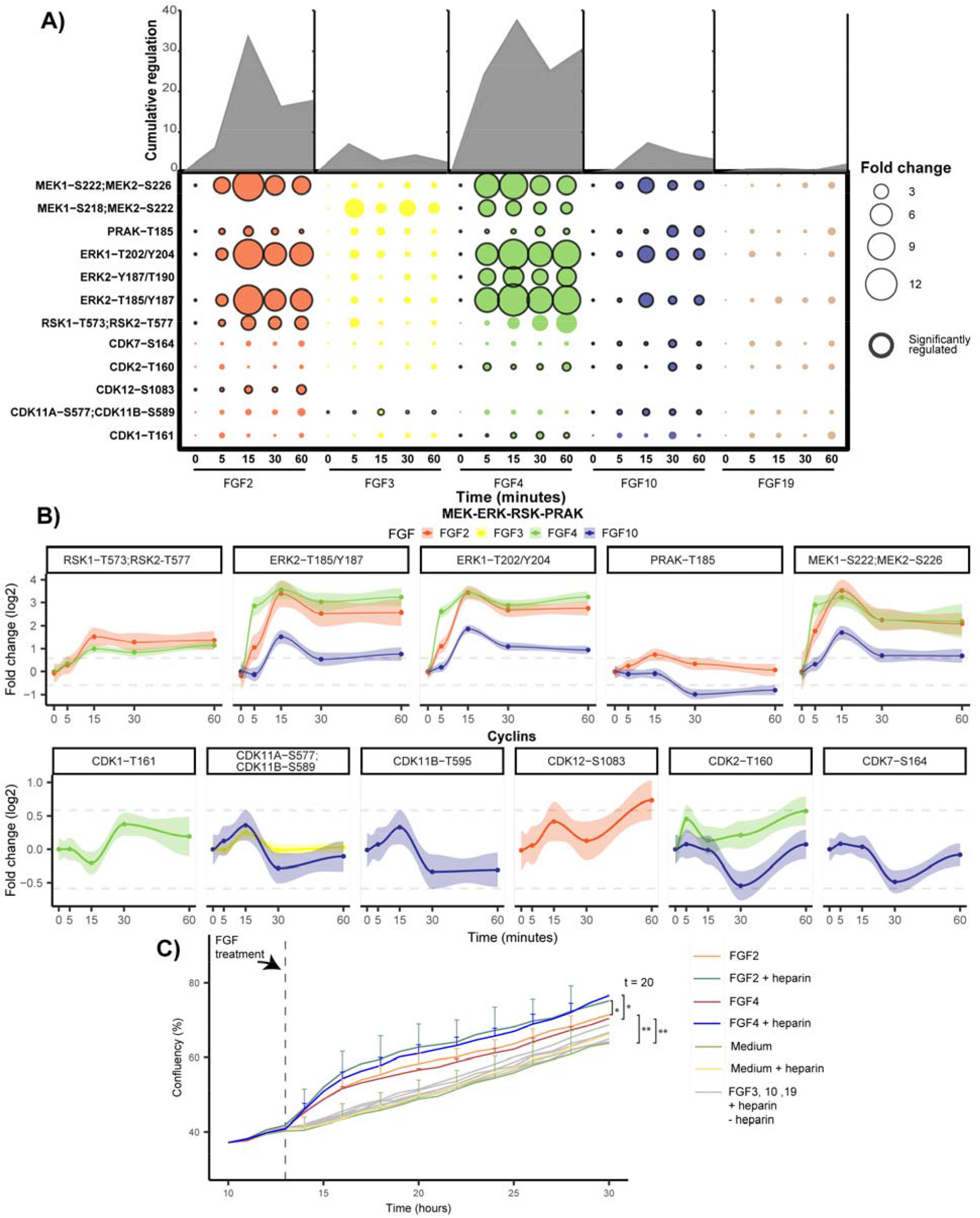
Regulation of kinases and cyclins implicated in the MAPK/ERK pathway. **A)** Measured phosphorylated peptides involved in the MAPK/ERK pathway that show significant regulation (ANOVA p < 0.05, independent triplicate measurements) following incubation with at least one of the tested FGFs. Black-edged circles represent significant phosphosites. The area of the circle represents cumulative regulation in fold change of the measured phosphopeptides measured in triplicate over 5 subsequent time points. The dynamic phosphorylation of the sites is color-coded by the FGF ligand used, and only plots are depicted when the ligand changed the phosphorylation at that site significantly. Grey lines represent a 1.5 fold-change, and 90% confidence intervals are presented per phosphopeptide. **C)** Influence of incubation with the FGF ligand and heparin on cellular growth. Growth curves of MCF-7 cells, incubated with 50ng/mL of each of the tested FGF ligands with or without 5µg/mL of heparin. The confluency percentage was taken as a readout to analyze cell growth and plotted (data was acquired in triplicate showing the standard deviations per time point and ligand used).

Investigating the kinases involved in the MAPK/ERK pathway after either FGF2, FGF4, or FGF10 treatment, showed rapid and high regulation of especially the main signaling hub of the MAPK/ERK pathway, namely MEK (MEK1 and MEK2) and ERK (ERK1 and ERK2) (**Figure 2B**) (Lavoie et al., 2020). FGF2 and FGF4 treatment resulted in an >10-fold increase of MEK and ERK activating phosphorylation and FGF10 an >2-fold increase. Notably, MEK and ERK activation was about 10-fold higher compared to the other kinases in the MAPK/ERK pathway, supporting their central role as signaling hub (**Figure 2B**). MEK and ERK dynamics per FGF treatment showed high correlation, displaying direct regulation of ERK as the target of MEK. However, MEK and ERK signaling dynamics showed a lower correlation between different FGF stimulations, also in the case of strong activation via FGF2 and FGF4. FGF4 treatment resulted in fast activation peaking at 15 min and showing an additional increase after the 30-minute time-point. FGF2 treatment on the other hand showed slightly slower activation with also the maximum at 15 minutes that plateaus from 30 minutes onwards (**Figure 2B**). This suggests differential MAPK/ERK pathway regulation.

Even though FGF2, FGF4, and FGF10 all activated MEK and ERK, all ligands resulted in unique downstream activation, which could either be the result of different activation mechanisms and feedback loops or due to different activation dynamics of the same pathway (Aoki et al., 2013; Raina et al., 2022). After FGF2 and FGF4 treatment, but not FGF10, MEK and ERK dynamics highly correlated with RSK1 and RSK2 dynamics, which are regulators of cell proliferation and cell survival (**Figure 2B**) (Anjum & Blenis, 2008; Houles & Roux, 2018; Romeo et al., 2012). FGF2 incubation resulted in the activation of CDK12 and a transient 1.5-fold increase in activating phosphorylation of PRAK, of which the role in the context of FGF has remained elusive (Maher, 1999; New, 1998). FGF4 incubation resulted in the activation of CDK1 and CDK2. Interestingly, CDK activation dynamics are relatively modest, with a maximum increase in activating phosphorylation of 60% (**Figure 2B**). Uniquely, FGF10 treatment did not activate kinases downstream of ERK but inactivated CDK2, CDK7, CDK11a, CDK11b, and PRAK. Inactivation of these kinases occurred concurrently after 30 minutes, which may originate from a negative feedback loop (Kuo et al., 2014). FGF10 may initiate this feedback loop by recycling its receptor FGFR2b to the cell membrane, or FGFR2b intracellular transport may expose the receptor to the substrates responsible for the feedback loop (Smith et al., 2021). Notably, only FGF10 showed sustained PRAK inactivation, which has been associated with decreased tumor progression (P. Sun et al., 2007; Y. Wang et al., 2021).

FGF3 and FGF19 have been described to activate the MAPK/ERK pathway in a subset of cell lines through FGFR4 activation (Desnoyers et al., 2008; Shi et al., 2009; Shinya et al., 2001; Teng et al., 2018, p. 201). In contrast, in our dataset, we did not observe any activation of the MAPK/ERK pathway after FGF3 and FGF19 stimulations, although, in our proteome profiles of MCF-7 cells after 24 hours of incubation with different FGFs, we did clearly identify the FGFR4 receptor.

In the context of FGF stimulation, MAPK/ERK pathway activation is considered to drive cell growth and increase tumor progression (Koledova et al., 2019; Lovicu & McAvoy, 2001; Y. Sun et al., 2017). To verify whether cell growth was indeed induced in our experimental conditions, we monitored cell growth after FGF stimulations using an IncuCyte ZOOM™. Only after stimulation with FGF2 and FGF4 we temporarily observed significantly increased cell growth (two-tailed t-test, p < 0.05) (**Figure 2C**). This finding was in line with the high MAPK/ERK pathway activation quantified in FGF2 and FGF4-stimulated cells. FGF10 stimulation did not substantially increase cell growth even though the MAPK/ERK pathway was activated. This suggests that a signaling threshold must be reached to activate proliferation or that alternative signaling is required for cell growth. Notably, adding heparin significantly increased the proliferation rate of FGF2 and FGF4-treated MCF-7 cells, while only adding heparin did not increase cell proliferation (**Figure 2C**).

### Consistent down-regulation of the PI3K and PKA pathway

Next, we examined the PI3K and PKA pathways. In our analysis, incubation with each of the tested FGFs, except FGF3, resulted in the significant inactivation of the PI3K and PKA pathways (**Figure 3A**). The PKA pathway is not commonly described to be regulated by FGFs. However, we quantified the change in phosphorylation of the upstream regulator PDPK1, which directly regulates PKA activity by phosphorylating Thr-197 (Cauthron et al., 1998). All measured kinases involved in the PKA pathway highly correlated with PDPK1 dynamics for all FGF stimulations in our dataset, revealing possible crosstalk between the PI3K and the PKA pathway. All tested FGFs, except for FGF3, resulted in similar inactivation of PDPK1 and the PKA pathway kinases PKA, GSK3A, and MARK kinases (**Figure 3B**). Inactivation was consistent but modest. The most significant decrease was a 2-fold decrease on two phosphorylated sites in the activation loop of MARK1, MARK2, and MARK3, respectively (**Figure S3**). Notably, no relation has been described between MARK kinases and FGF signaling up to this day. MARK kinases control cell polarity by regulating microtubules, and reduced MARK kinase activity has been linked to EMT, which is in line with the EMT-inducing effects of FGFs (Drewes et al., 1997; Sonntag et al., 2017).

**Figure 3.**
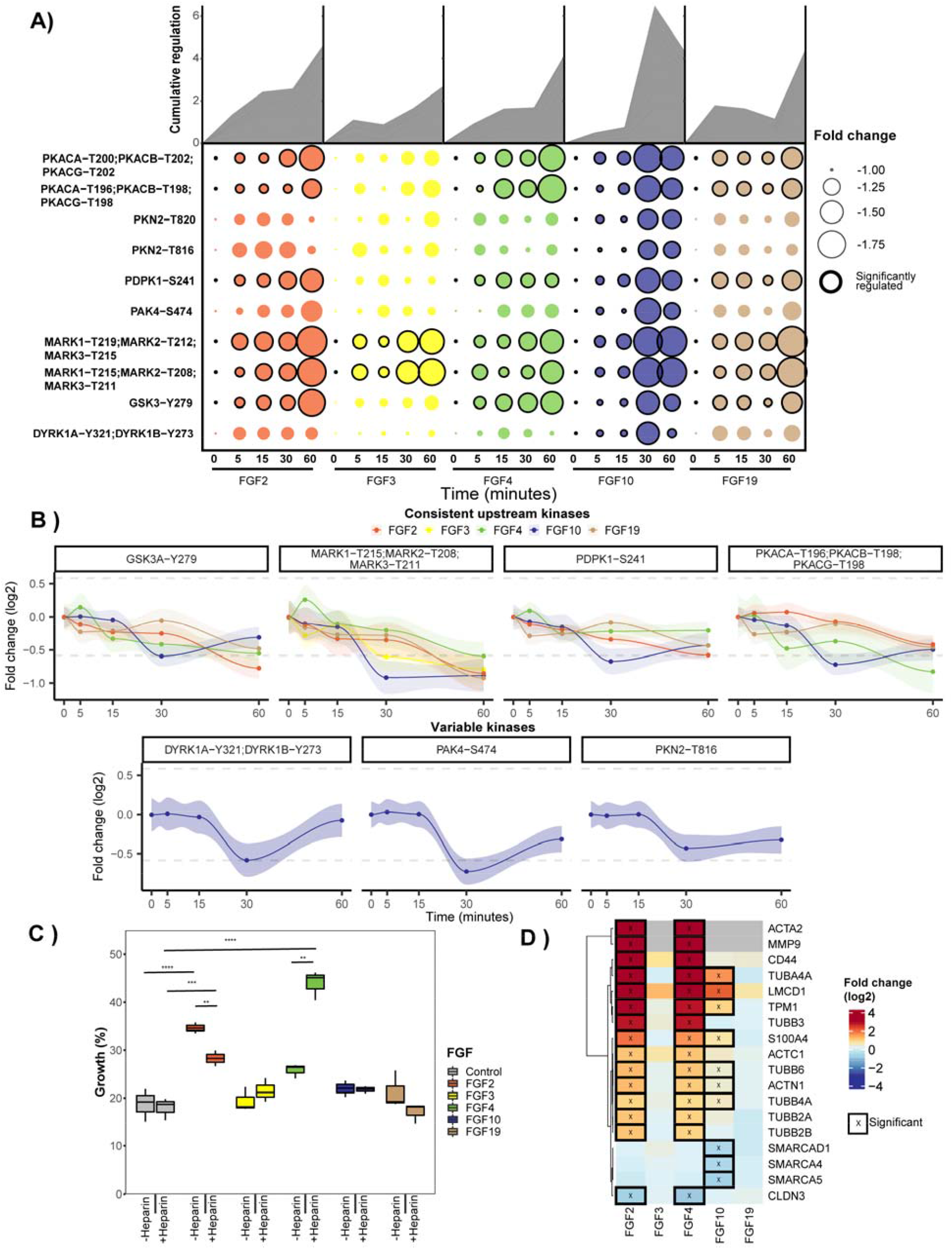
Regulation of kinases implicated in the PKA and PI3K pathways. **A)** All significantly regulated phosphopeptides of the PI3K pathway are represented as the mean across triplicates (p < 0.05 ANOVA). Cumulative absolute regulation in fold change is represented in the area plot to show overall pathway regulation. **B)** Regulation of significant (ANOVA p < 0.05) changing phosphopeptides are plotted from the PI3K pathway. Grey lines represent a 1.5 fold-change and 90% confidence intervals are presented per phosphopeptide. **C)** MCF-7 cells were subjected to a scratch wound assay, and after 24 hours percentage closure of the scratch was measured. The boxplots represent triplicate measurements of FGF-stimulated cells with or without 5 µg/µL heparin. A two-sided t-test was used to validate the significance. **D)** Proteins associated with EMT were extracted from the shotgun dataset. Significantly regulated proteins are displayed using an asterisk (FDR < 0.05).

Besides the similar PI3K and PKA pathway regulation by FGF2, FGF4, FGF10, and FGF19, solely FGF10 stimulation led to a decrease in phosphorylation of the downstream kinases PAK4, DYRK1A, DYRK1B, and PKN (**Figure 2B**). This reveals broad FGF10-induced negative regulatory mechanisms. Notably, the inhibited PAK4, which plays a role in cell adhesion, can be regulated via ERK and the PI3K pathway (Qu et al., 2001; Ramos-Alvarez & Jensen, 2018; Thillai et al., 2017; Won et al., 2019). The observed kinase activity dynamics of PAK4 strongly correlated with the rest of the PI3K pathway while opposing the MAPK/ERK pathway activity dynamics. This observation suggests that PAK4 is linked to the PI3K pathway, not the MAPK/ERK pathway.

FGF2, FGF4, FGF10, and FGF19 have all been described to induce EMT, which is thought to be partially regulated via the regulation of the PI3K pathway and is regarded as a key process in regulating tumor metastasis (Katoh & Katoh, 2006; B. P. Zhou et al., 2004). This agrees with the targeted kinome data that revealed PI3K pathway inactivation for these FGFs. To investigate whether the tested FGFs induced EMT, we next performed a wound-healing assay that assays cell migration capabilities, which is a key process in EMT (Grada et al., 2017; Yan et al., 2010). These assays revealed that, only FGF2 and FGF4 showed a significant increase in wound-healing capacity of 15 and 5% compared to unstimulated MCF-7 cells, respectively (**Figure 3C**). Interestingly, with the addition of heparin, this dampened to 10% for FGF2 and increased to 40% for FGF4, revealing a modest role for heparin in regulating EMT. To find further support for FGF-induced EMT, we extracted proteins from the EMTome database associated with EMT, specifically focusing on proteins that directly trigger EMT or are key markers for EMT (Vasaikar et al., 2021). In their proteomic profiles (**Table S3**), FGF2 and FGF4 stimulations showed an identical profile of 15 EMT-associated proteins significantly regulated after 24 hours (**Figure 3D**), supporting an EMT-like phenotype downstream of FGF2 and FGF4. FGF10 stimulation resulted in less pronounced expression changes in 7 of the 15 observed EMT proteins, in part confirming the role of FGF10 in inducing EMT, whereas FGF3 and FGF19 showed no significant expression changes in EMT-related proteins (Brabletz et al., 2018). This is further supported by GSEA analysis of the hallmarks of EMT as provided by MSigDB, which in our proteome data are only significantly upregulated after FGF2 and FGF4 treatment (**Figure S4**) (Subramanian et al., 2005). Altogether, these findings show FGF2, FGF4, FGF10, FGF19 all inactivated the PI3K pathway. However, only FGF2 and FGF4-treatment resulted in increased wound healing capacities and an EMT-like phenotype on proteome level, FGF10 treatment only resulted in a more EMT-like phenotype on proteome level, and FGF19 did not show either. Besides PI3K inactivation, further mechanisms must thus be regulated to induce EMT.

### Undistinguished PLCγ signaling along the FGF-FGFR axis

Next, we explored the measured activity profiles of the kinases within the PLCγ pathway. The PLCγ pathway is relatively understudied in the context of FGFR stimulation and regulates specialized functions (Kim et al., 2003; Lima et al., 2009; Mohammadi et al., 1991; Niger et al., 2010; Ranieri et al., 2020; Szybowska et al., 2021). In our current study, PKD1, PKD2, PKD3, PKCδ, and PKCγ showed significant regulation when incubated with at least one of the tested FGFs (**Figure S5**). Kinase activation dynamics were non-linear, hinting at the presence of multiple feedback loops (Kuo et al., 2014). Moreover, kinases in the PLCγ pathway showed a relatively low correlation in their activation dynamics, and all tested FGFs showed distinct kinase regulation (**Figure S5**).

Indeed, only FGF2 transiently activated PKCα/β/γ by activating phosphorylation Thr-514, yet no other PLCγ pathway kinases were regulated (Kelher et al., 2017). FGF4, FGF10, and FGF19 all activated PKD1 and PKD3, whereas FGF4 and FGF10 also showed the inactivation of PKCδ and PKD2 or only PKCδ, respectively.

### Distinct FGF ligands induce distinct and diverse temporal dynamics in phospho-signaling

Not only does FGF specificity to the various FGFRs determine the biological outcome, but also the affinity for the various FGFRs is crucial. In RTKs biology, it is known that ligands with high affinity to the receptor can lead to fast, transient activation, while lower affinity ligands, binding to the same receptor, lead to a slower sustained activation, resulting in a different biological outcome (Huang et al., 2017; Kiyatkin et al., 2020). To evaluate whether each FGF differentially regulated signaling dynamics, OmniPath was used to construct biological networks in which kinases are ordered based on the initial time point when regulation was observed (**Figure 5**) (Türei et al., 2016).

**Figure 5.**
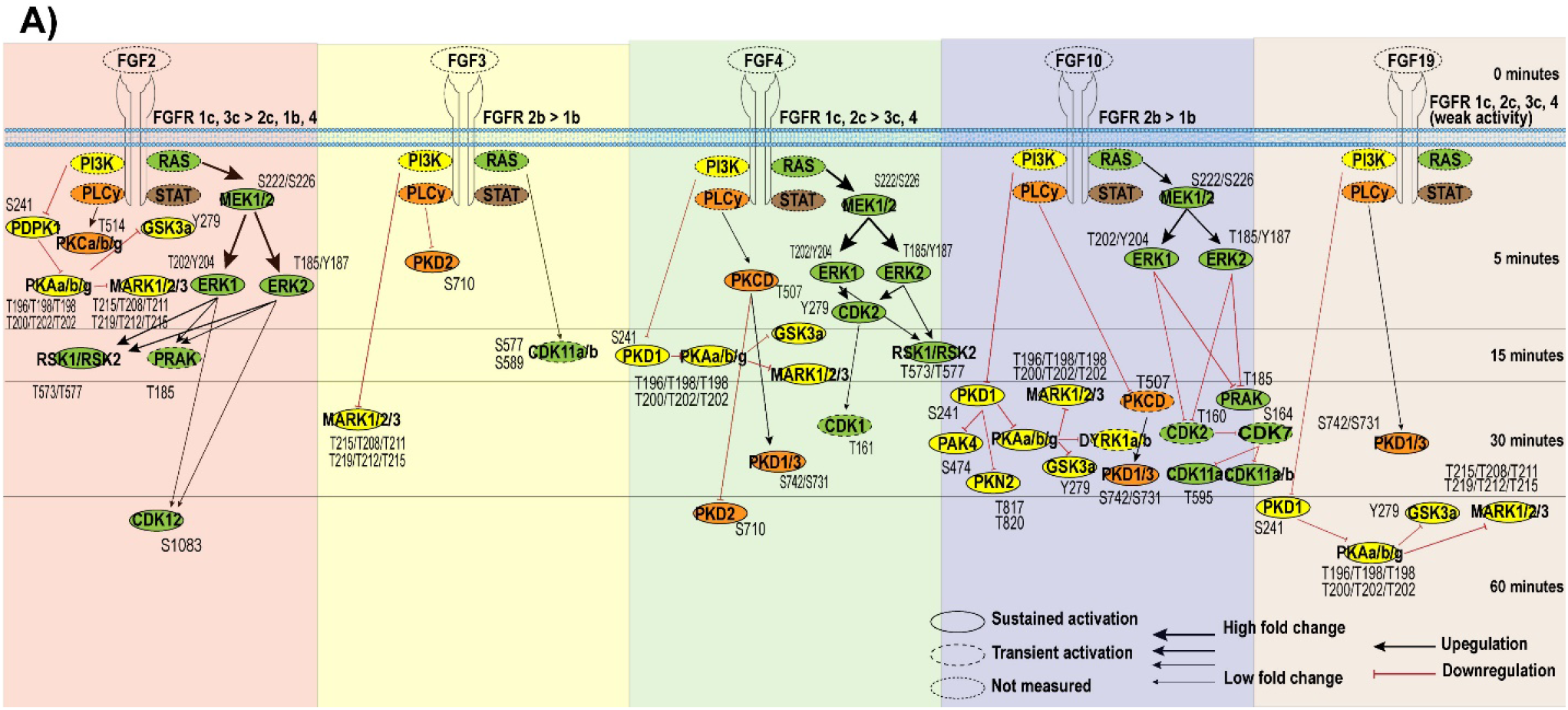
Temporal kinome dynamics following FGF treatment. **A)** MCF-7 cells were treated with either FGF2, FGF3, FGF4, FGF10, or FGF19, together with heparin. The resulting temporal kinome dynamics were quantified using the targeted activation loop assay and normalized to the t=0 time point. The presented kinome signaling dynamics are separated per FGF used for stimulation, and kinases are ordered based on significant initial activation (ANOVA + Tukey’s range test, p < 0.05, biological triplicates). Black and red arrows indicate whether measured kinase activity increased or decreased over time, respectively. The fold change compared to the t=0 time point is represented by the thickness of the arrows.

Indeed, in our data, each of the tested FGFs did lead to distinct timing of initial pathway regulation. FGF2 induced a fast initial activation within 5 minutes of all measured pathways. This is expected as FGF2 binds to most FGFRs with high affinity (Ornitz et al., 1996). Although FGF4 binds the same FGFR subset as FGF2, except for FGFR1b, it does so with different affinity. As a potential consequence, and in contrast to FGF2, FGF4 stimulation inactivated the PI3K and PKA pathways in our experiments only after 15 minutes (**Figure 5**). FGF10 stimulation activated the MAPK/ERK pathway within 5 minutes, similar to FGF2 and FGF4. However, this was followed by a strong downregulation after 30 minutes of more downstream targets. Last, FGF3 and FGF19 resulted in relatively slow (and modest) activation only 30 minutes after stimulation (**Figure 5**).

### Logic-based dynamic modeling validates the known FGF pathways but also identifies putative new players

Pathway models such as in Figure 5 are based on existing knowledge and are thus inherently biased towards well-characterized pathways. Therefore, validation of the biological model is needed to identify either missing or inaccurate connections between kinases or missing signaling nodes. To verify our biological model, predict signaling dynamics between kinases, and find possible gaps, we used a dynamic mechanistic model based on logic-based ordinary differential equations (Morris et al., 2010). First, a prior knowledge network (PKN) was built using information available via OmniPath using only kinases quantified in all FGF stimulations (**Figure 5**) (Türei et al., 2016). Next, the logistic-based ordinal differentiations were calculated using the quantitative longitudinal kinase activity data of all FGF stimulations together. For each node, a speed factor (τ) was calculated to represent the responsiveness of a kinase’s activation to upstream kinases activation (Wittmann et al., 2009). Low values indicate a slow transfer of activation from kinases’ upstream activators. For each node, also an edge-specific transmission parameter (k) was calculated, which represents the quantitative signal that is transferred between kinases (Wittmann et al., 2009). High values of the non-linear k parameters indicate that relatively little quantitative signal is transferred via the edge. To evaluate the quality of the predicted τ and k values, Pearson’s r and the RMSE of all the quantitative kinome values in the model were assessed and compared to the RMSE between biological replicates (**Figure 6A** and **B**). The RMSE of the model (0.18) is almost as low as the RMSE observed between the biological replicates (0.1). The model thus successfully predicts most of the kinase activity, with a small error likely due to unknown entries in the PKN.

**Figure 6.**
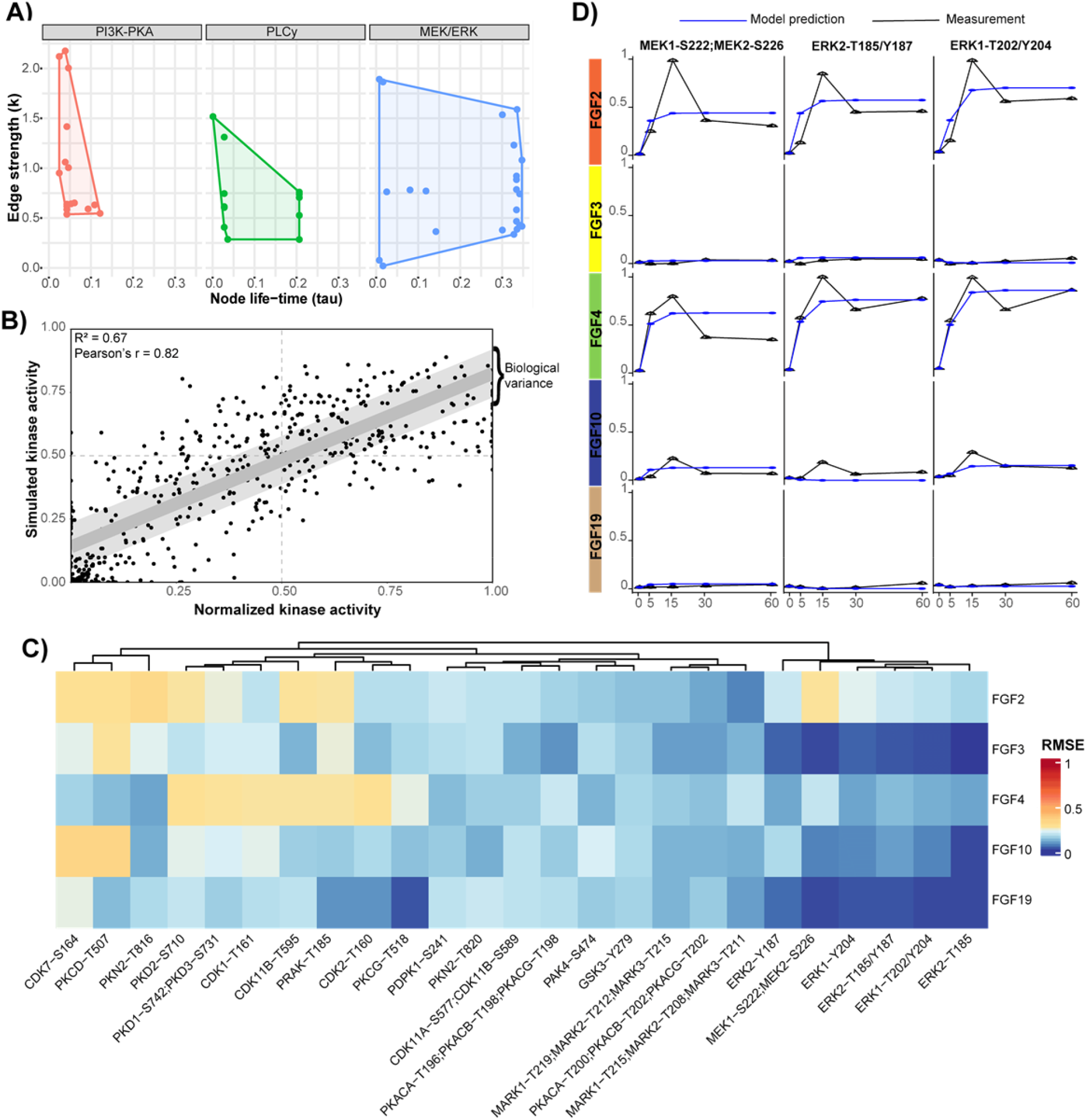
Logic-based dynamic modeling reveals unknowns in FGF-induced signaling. **A)** Logic-based dynamic modeling was used to predict a speed factor (τ) and a transmission parameter (k) for the kinases across the PI3K, PKA, PLCγ, and MAPK/ERK pathways. These represent the signal transduction speed and the quantitative signal transferred between kinases, respectively. **B)** The root mean squared error (RMSE) of the predicted values by logic-based dynamic modeling and the measured values by the targeted kinome loops assay. The values were normalized using the 99% interquartile range. The light grey area represents the biological variation in the measurements. The dots represented are the mean values of the replicates and all time points. **C)** Mean RMSE values for the measured vs. predicted kinase activity values. The modeling was performed using identical networks, meaning downstream kinase-kinase relations constitute the same predictive k and tau values. Therefore, predictive downstream errors may indicate differential regulation between FGFs. **D)** Line plots of the measured and predicted kinase activity using the function with the lowest error across all FGF stimulations. Before the logic-based dynamic modeling, the average of the quantified kinome values was taken (biological triplicates) and normalized using the 99% interquartile range. The blue line represents the model prediction, and the black line represents the quantified kinase activity using the targeted kinome assay.

To explore these unknowns in the PKN, the RMSE of individual phosphopeptides was evaluated (**Figure 6C**). High RMSE suggests that the model is insufficient to predict a kinase activation state, which results from missing or erroneous connections between nodes in the network. Therefore, a high predictive error can be used to find novel biological connections or nodes. The model showed no highly contradictive prediction errors for single kinases (RMSE error > 0.5), which occurs when activation of one kinase leads to activation of the next kinase, but inactivation is measured. However, some kinases showed errors that were higher than the biological variance.

Kinases with a relatively high error are part of the PLCγ and MAPK/ERK pathways. Error in kinases regulated by the PLCγ pathway is expected due to the low pathway coverage (**Figure 5**). Surprising, however, is the substantial error in MEK activity prediction after FGF2 stimulation (**Figure 6C**). The model failed to predict the fast activation of MEK and ERK, and did not incorporate the oscillatory patterns typical for feedback loops (**Figure 6D**). Further, following MEK-ERK signaling downstream, all measured CDKs, including CDK1, CDK2, CDK7, CDK11a, and CDK11b, show a relatively high predictive error. This suggests differential MEK-ERK signaling to their downstream effectors. We, therefore, hypothesized that the error in MEK-ERK-CDK signaling is indicative of unknown links between kinases or missing nodes in the current model. We will focus on this more in the next section.

### Modeling guided analysis unveils differential FGF signaling

With the aim to explain the predictive modeling errors for MEK, ERK, and CDKs, we expanded the model using manually curated literature mining, shotgun phosphoproteomics analysis of FGF-stimulated MCF-7 cells (**Table S4**), and our targeted kinome data. Significantly regulated proteins were used to construct a more refined pathway (**Figure 7A**) (Gotoh, 2008; Hadari et al., 1998; Yang et al., 2006).

**Figure 7.**
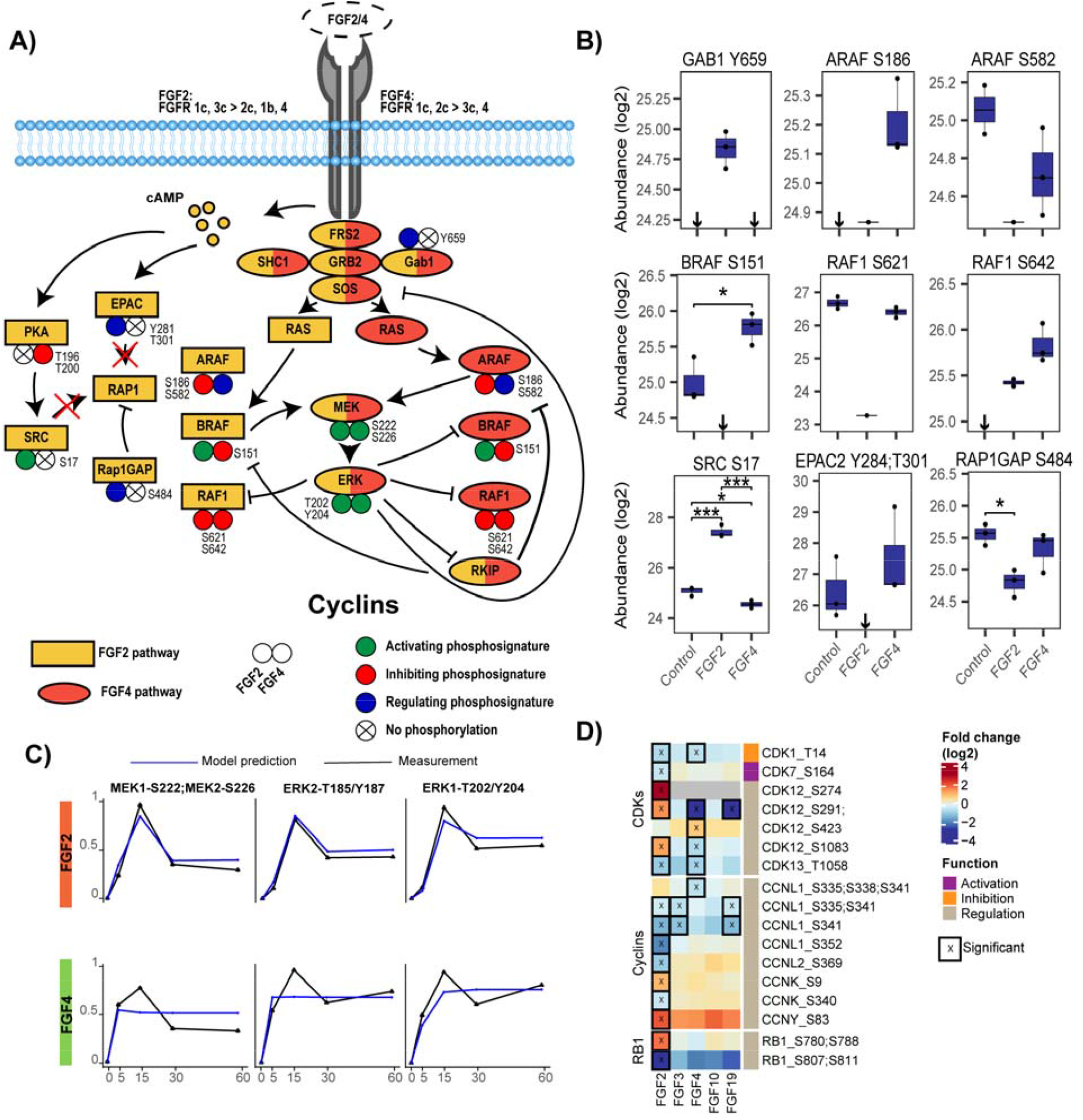
Regulation of RAF family kinases modulates ERK signaling. **A)** Mapping of phosphorylations of proteins involved in ERK activation shows tight regulation of the RAF kinase family members. The regulation does occur in the RAP1 activation pathway of ERK, yet no RAP1 activation was measured, suggesting this does not contribute to ERK activation. **B)** Quantified peptide abundances corresponding to figure 7A. Significance is depicted using * (p < 0.05) or *** (p < 0.001) using a two-sided t-test (biological triplicates). If all values are below the detection limit, this is shown using a ↓. Abundances are acquired using shotgun phosphoproteomics after 60-minute stimulation with the different FGFs. **C)** Line plots of the measured kinase activity and the predicted kinase activity using the function with the lowest error across all FGF stimulations. The PKN used is the updated biological pathways, also presented in figure 7A. The average of biological triplicates was taken and normalized using the 99% interquartile range. Model predictions are shown in blue, and quantified kinase activity is shown in black. **D)** Phosphorylation of cell cycle regulating proteins from the phosphoproteomics data. Significant regulated sites are displayed (two-sided t-test, p < 0.05, biological triplicates). Scores represent log2 fold changes.

A potential missing link came from the phosphoproteomics data that suggested a RAP1 activation signature exclusively for FGF2. RAP1 is an alternative activator of MEK-ERK, whose activators include EPAC2 and SRC, and its main negative regulator is RAP1gap (**Figure 7A**) (Looi et al., 2020; Schmitt & Stork, 2002; Stokman et al., 2014). Uniquely, FGF2 treatment abolished the signal of Tyr-284 and Thr-301 phosphorylation of EPAC2, which is important for EPAC2 membrane localization. Moreover, FGF2 treatment increased activating phosphorylation Ser-17 of SRC about 4-fold and resulted in a 1.6-fold increase in regulating phosphorylation Ser-484 on RAP1gap (Daumke et al., 2004; de Rooij et al., 2000; Fukuyama et al., 2005, 2006; Rehmann et al., 2003, 2006). These phosphorylations highlight possible RAP1 activation. Therefore, we conducted a RAP1 activity assay. However, this RAP1 activation assay showed no significant RAP1 activation in all tested ligands (**Figure S6**). From these data, we concluded that although pathways commonly involved in RAP1 activation were regulated, RAP1 was not activated and thus was not the cause of differential MEK-ERK dynamics.

Next, we compared FGF2 and FGF4-induced signaling along a more detailed RAS-RAF-MEK-ERK signaling axis (**Figure 7A** and **B**). FGF2 and FGF4 treatment resulted in fine-tuned and distinct regulation along this signaling axis, especially of the RAF family members (ARAF, BRAF, RAF1) that coordinate MEK-ERK activation (Maurer et al., 2011). Solely FGF2 treatment enabled BRAF activity by abolishing the signal of the inhibitory ERK target site Ser-151 on BRAF (**Figure 7B**) (Marquette et al., 2011). Moreover, FGF2 treatment resulted in reduced activity of ARAF following significant downregulation of Ser-582 phosphorylation, which is needed for 14-3-3 binding to increase the activity of ARAF (Baljuls et al., 2008). FGF2 also resulted in a reduced active state of RAF1 implied by an 8-fold lower signal of Ser-621 phosphorylation, necessary for 14-3-3 activation, and by the negative feedback phosphorylation of Ser-642 by ERK on RAF1 (**Figure 7B**) (Dhillon et al., 2009; Zang et al., 2008) (Dougherty et al., 2005). Contrarily, FGF4 stimulation showed an activating signature for ARAF, indicated by the phosphorylation of the regulatory site Ser-186 on ARAF (Stuart et al., 2015). Further, FGF4 stimulation resulted in inhibitory phosphorylation on BRAF and RAF1, with a twofold increase in Ser-151 phosphorylation on BRAF, and a strong increase in Ser-642 phosphorylation on RAF1, which was absent in the control (**Figure 7B**). In conclusion, the RAF family members showed differential regulation as FGF2 treatment indicated BRAF-driven activation, while FGF4 treatment indicated ARAF-driven activation.

To further validate these signaling differences, we again applied logic-based dynamic modeling using the data from the targeted kinome assay. In the updated model, FGF2 signaling was directed via BRAF and FGF4 via ARAF. Moreover, to model the negative feedback loops, one negative feedback loop between ERK and the FGF activation of ARAF and BRAF was added, as well as a negative feedback loop from ERK to RKIP and from RKIP to ARAF and BRAF activation of MEK (Shin et al., 2009). The updated pathway showed improved modeling accuracy (**Figure 7C**). Especially the FGF2 signaling prediction now has high accuracy that follows the measured feedback loops, giving confidence to the predicted biological pathway. Prediction of FGF4 signaling dynamics was also improved over the initial model, with more accurately modeled activation dynamics, however, is not optimal yet (**Figure 7C**). Indeed, the updated model supports the two different modes of ERK activation downstream of FGF2 and FGF4, yet, alternative regulators need to be identified to fully explain FGF signaling dynamics.

Following ERK activation further downstream, we set out to use the shotgun phosphoproteomics data to confirm predictive errors for the CDKs and validate differential regulation downstream of ERK. Cumulatively, 17 different phosphorylated sites on proteins that regulate the cell cycle were quantified, including CDKs, cyclins that regulate CDK activity, and RB1, which are all central to cell cycle progression (**Figure 7D**) (Loog & Morgan, 2005; Valverde et al., 2022). FGF3, FGF10, and FGF19 showed little CDK regulation in our model, in line with the targeted kinome data and the modeling results. FGF2 and FGF4 showed distinct activation patterns of CDKs (**Figure 7D**), agreeing with the targeted kinome data and the modeling error. These distinct activation patterns confirm the predictive error of the dynamic model and show that FGF2 and FGF4 regulate cell cycle progression differently.

## Discussion

By investigating the FGF-induced dynamic kinome regulation using a targeted kinome assay, we quantified and compared the signaling responses of FGF2, FGF3, FGF4, FGF10, and FGF19. All FGF stimulations resulted in a unique biological response in MCF-7 cells, with FGF2 and FGF4 having the broadest kinome response, FGF10 having a moderate response, and FGF3 and FGF19 showing a modest response. We find complex activation mechanisms that initiate FGF signaling as biological responses upon FGF stimulation vary between cell lines, do not correlate with FGFR expression level and are influenced by heparin.

Looking at the FGFs in a breast cancer context, FGF-stimulated cells activate biological pathways that can contribute to the hallmarks of cancer (Hanahan & Weinberg, 2011; Xie et al., 2020). The MAPK/ERK pathway is thought to drive cell proliferation, and the PI3K pathway is believed to regulate EMT (Kunath et al., 2007). However, we find that simply activating these pathways does not per se lead to cell proliferation or EMT, respectively. Importantly, this irregularity between kinome or pathway activation and predicted biological outcome emphasizes the complexity of these processes and their incomplete understanding. FGF2 and FGF4 increased cell proliferation and EMT in MCF-7 cells. However, FGF3, FGF10, and FGF19 are reported to regulate cell proliferation and EMT but were not able to regulate these processes in our system. Additional signaling factors may be needed to sensitize or co-stimulate the cells for a more pronounced biological response (Desnoyers et al., 2008; W. Wang et al., 2015; Watson & Francavilla, 2018).

The quantification of dynamic kinase responses instead of single time points is highly advantageous for understanding FGF-stimulated signaling because these dynamics expose unknown signaling routes and improve the reliability of the predicted signaling network. Often biological networks are deduced from literature without proper validation. For this purpose, logic-based dynamic modeling provides a suitable solution. Logic-based dynamic modeling of the FGF stimulations resulted in an overall low network error implying feasible network predictions. Mainly the PLCγ pathway showed higher predictive errors due to a higher sparsity of the network, partly due to limited insights into PLCγ signaling in the FGF context. This highlights the importance of further studying PLCγ signaling to understand its functions in FGF signaling (Brewer et al., 2016).

The dynamic modeling highlighted differential and fine-tuned regulation of the MAPK/ERK pathway. Regulating phosphorylations of the RAF kinases indicate that FGF2 stimulation is directed via BRAF, while FGF4 stimulation is directed via ARAF. Literature on RAF kinase family regulation by FGFs is limited; however, understanding RAF regulation is essential because different RAF kinases perform different biological functions (Dumaz, 2011; Wellbrock et al., 2004). Moreover, understanding RAF signaling provides targeted insights that can be exploited to successfully deploy RAF specific inhibitors in various diseases, such as cancer (Saini et al., 2013). For example, Metzner et al., show that FGF-driven melanoma is, in some cases, sensitive to the BRAF inhibitor RG7204 (Metzner et al., 2011).

To conclude, this study highlights the differential signaling of FGFs and adopts existing logic-based dynamic modeling techniques to direct, strengthen, and increase the discovered biological knowledge.

## Acknowledgments

We thank Mara Diks and Suzan Thijssen (Utrecht University) for their guidance with the qPCR experiments. We thank Jennifer Haworth and Joseph Parsons (CF team) for providing qPCR sequences. This work has been supported by EPIC-XS, project number 823839, funded by the Horizon 2020 programme of the European Union and the NWO funded Netherlands Proteomics Centre through the National Road Map for Large-scale Infrastructures program X-Omics, Project 184.034.019. Research in CF lab is supported by the Wellcome Trust (107636/Z/15/Z and 107636/Z/15/A), the Biotechnology and Biological Sciences Research Council (BB/R015864/1), and Medical Research Council (MR/T016043/1).

## Author contributions

**Tim S. Veth**: Conceptualization; Data curation; Formal analysis; Investigation; Resources; Software; Visualization; Writing – original draft. **Chiara Francavilla**: Conceptualization; Writing – review & editing. **Albert J.R. Heck**: Conceptualization; Funding acquisition; Resources; Writing – review & editing. **Maarten A.F.M. Altelaar**: Conceptualization; Funding acquisition; Resources; Writing – review & editing.

## Data availability

Data are available via ProteomeXchange with identifier PXD038836 : (Username; Password: CCMgJt8J)

Data are available via ProteomeXchange with identifier PXD038808: (Username; Password: mYXS1PfR)

Data are available via ProteomeXchange with identifier PXD039005: (Username; Password: AIheltWY)

The used R scripts are available at: https://github.com/TVeth/FGF_Veth_2023

## Supplemental information

**Supplementary Figure 1:**
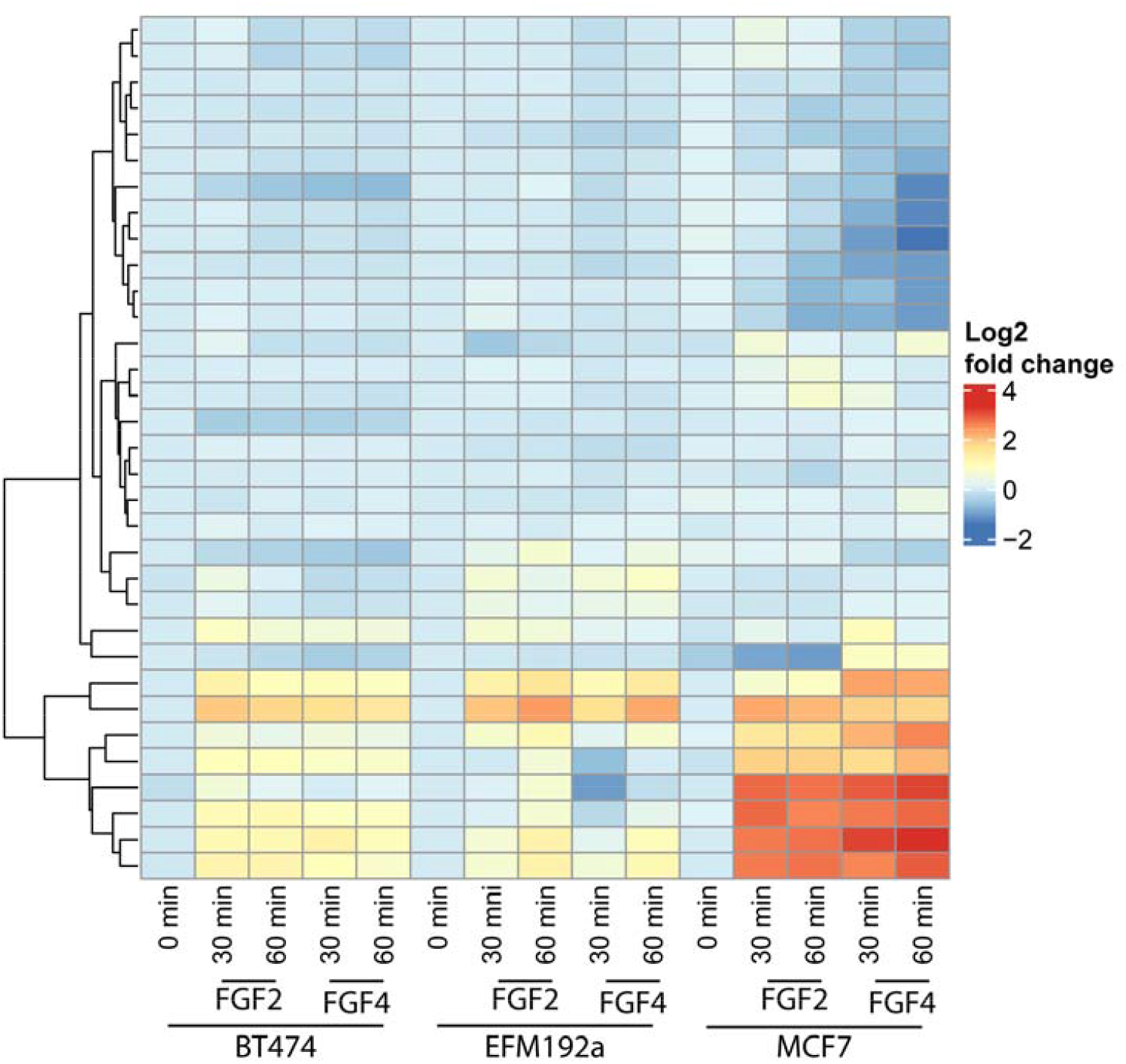
Kinome activity after FGF stimulation in breast cancer cells. MCF-7, BT-474, and EFM-192a cells were stimulated with 50ng/mL of either FGF2 or FGF4 supplemented with 5µg/mL of heparin. After 0, 30, and 60 minutes, the cells were harvested and subjected to measurement using the targeted kinome assay. The heatmap shows the quantified activation-determining phosphorylated sites on the kinases (biological triplicates). Only significantly changing values are shown (ANOVA p < 0.05).

**Supplementary Figure 2:**
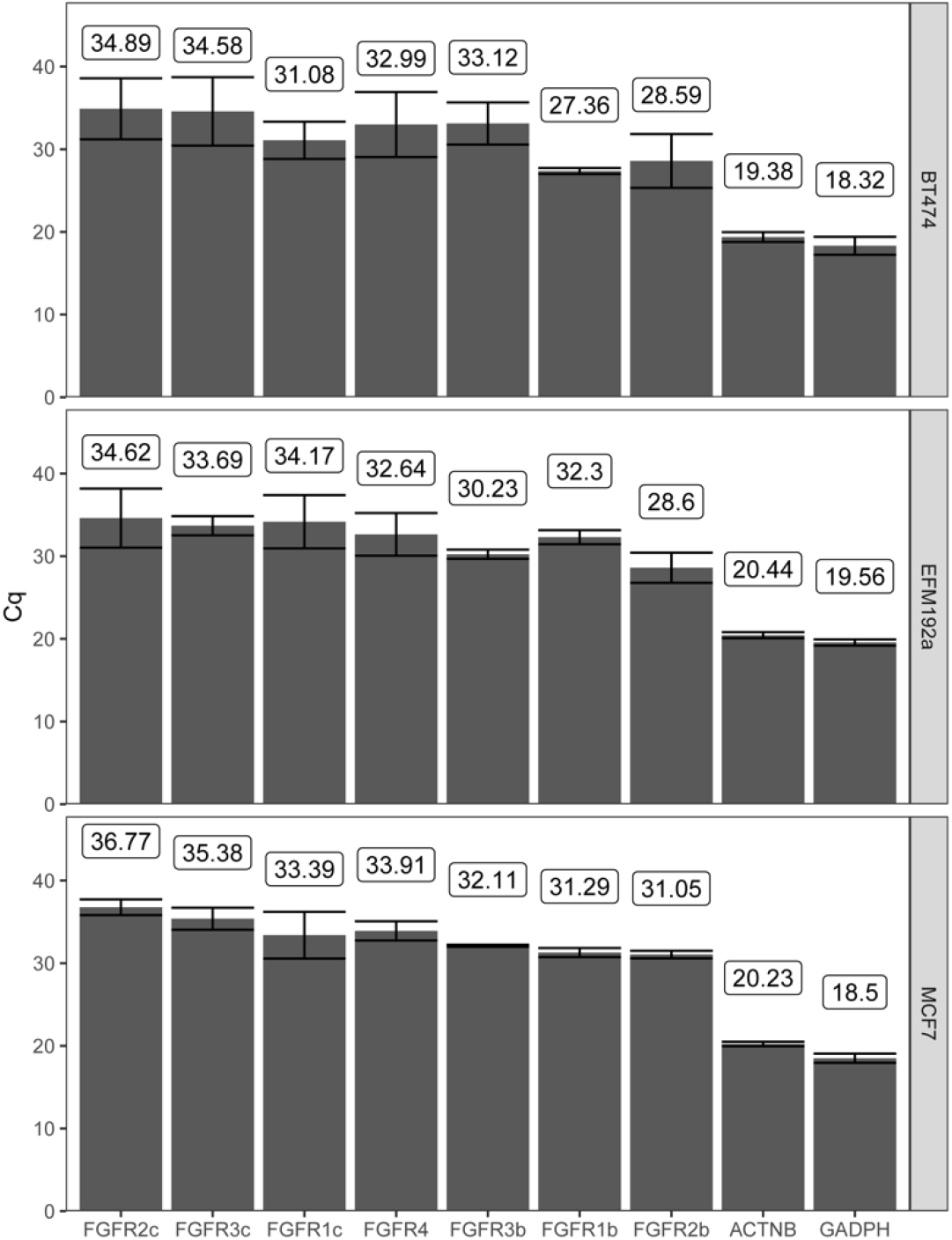
qPCR quantification of FGFR abundance. FGFR expression levels were quantified using qPCR in MCF-7, BT-474, and EFM-192a cells using FGFR subtype-specific primers (triplicate measurements) (**Supplementary table 5**). Beta-actin and Glyceraldehyde-3-phosphate dehydrogenase was quantified to enable normalization across cell types. Reported values are the quantitation cycles (Cq) that negatively correlates with the RNA expression levels.

**Supplementary Figure 3:**
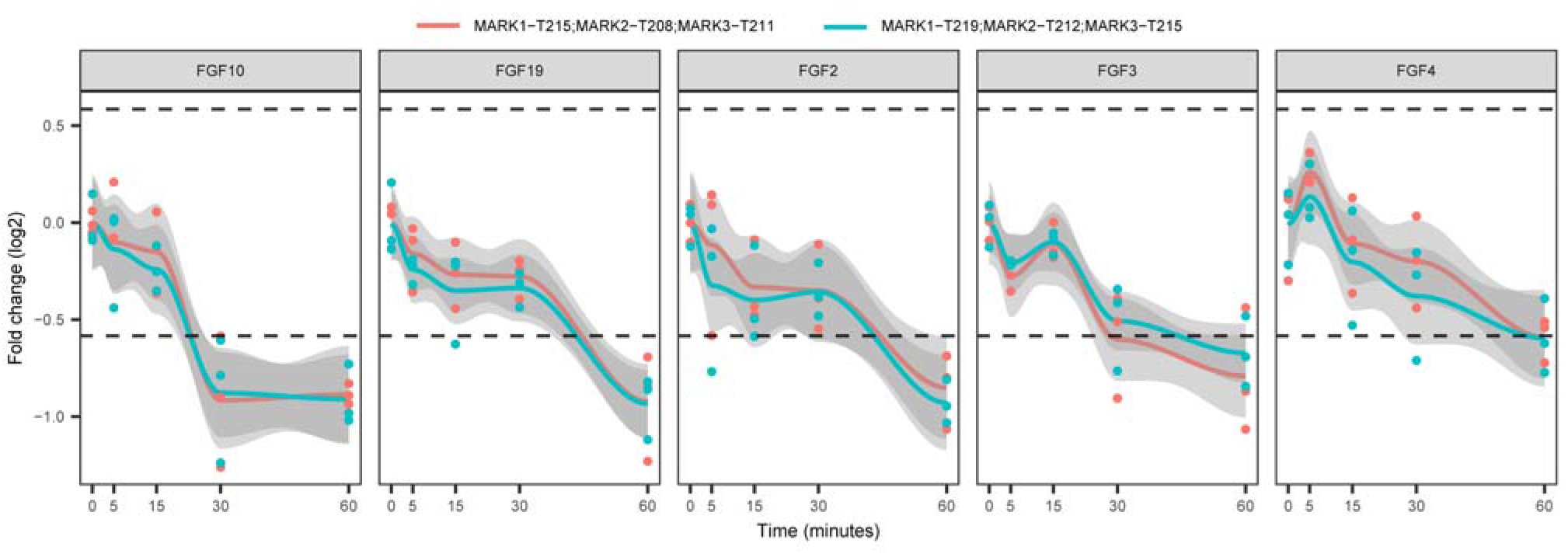
FGF induced MARK kinase regulation. Activity-determining phosphorylation in the activation loop of MARK1, MARK2, and MARK3 was quantified in MCF-7 treated with different FGF ligands. Line plots show these quantified phosphorylated sites (biological triplicates). Values are represented in log2 and the 1.5 fold-change is represented using the dashed line.

**Supplementary figure 4:**
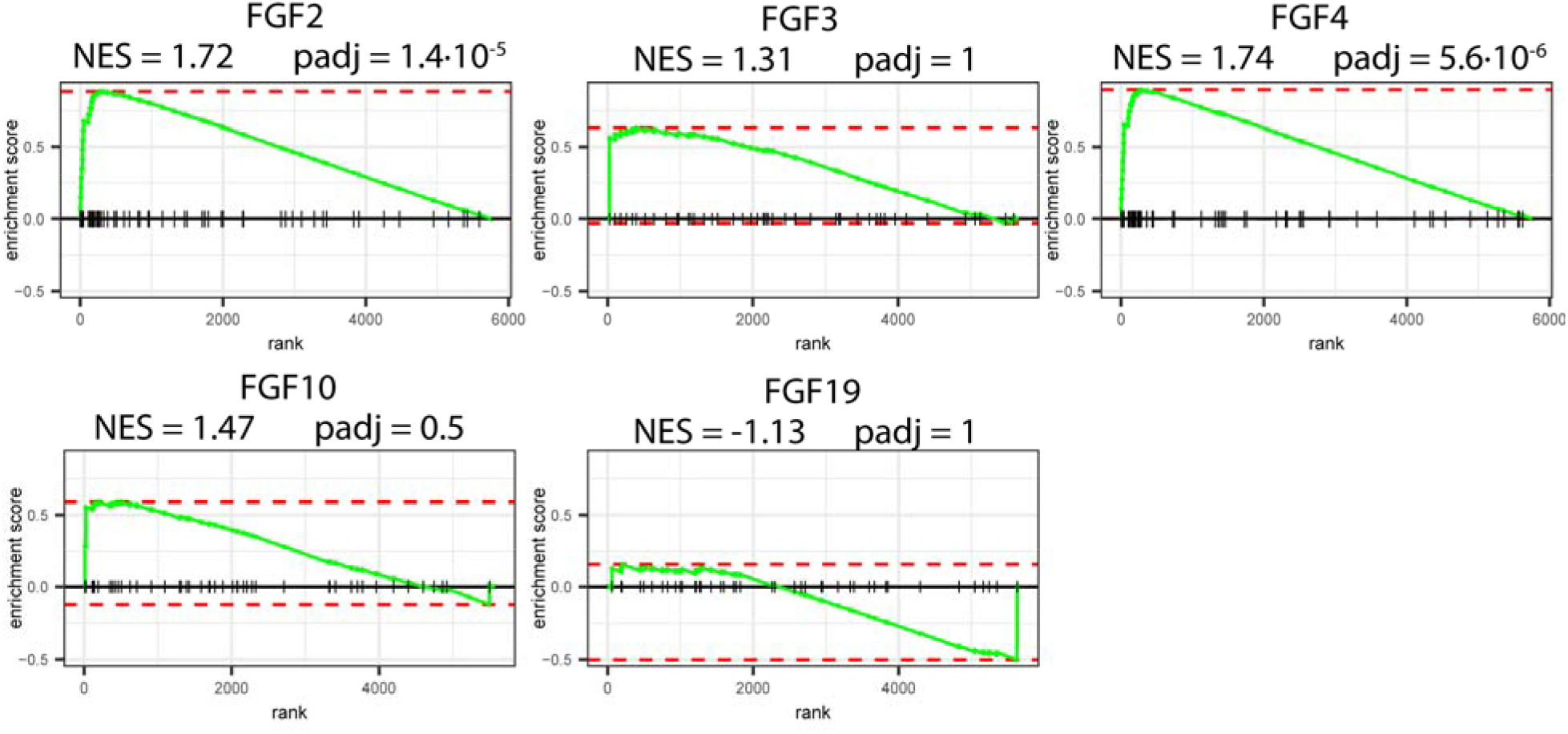
proteome derived EMT signature. MCF-7 cells treated were treated for 24 hours with the FGF ligands and their proteomes quantified. Subsequently, GSEA analysis on these proteomes (biological triplicates) was performed using the MsigDB signature “HALLMARK_EPITHELIAL_MESENCHYMAL_TRANSITION”. The normalized enrichment score (NES) along with the adjusted p-value is reported.

**Supplementary Figure 5:**
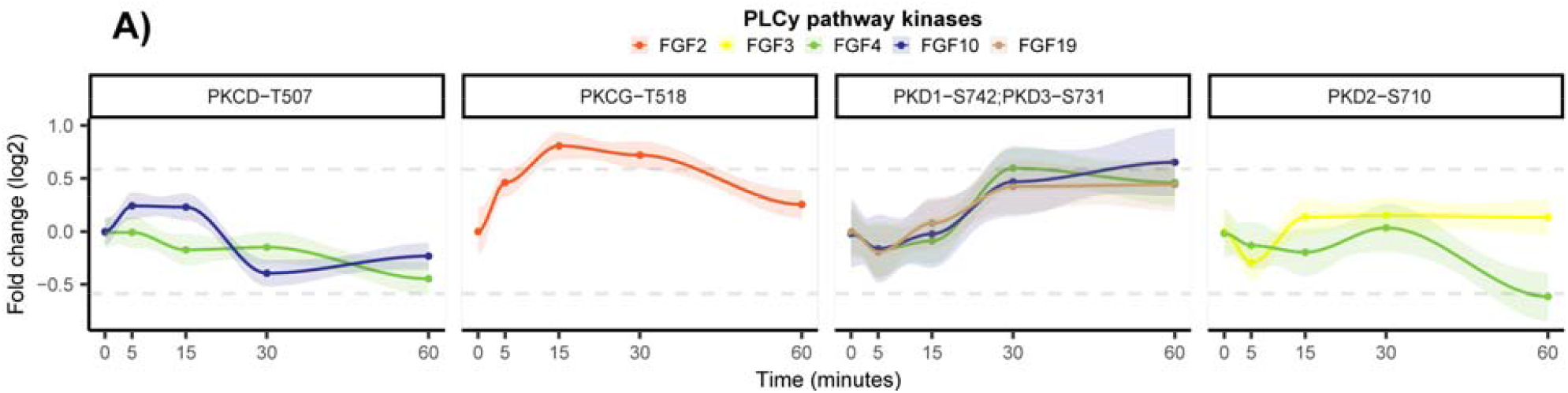
Regulation of kinases implicated in PLCγ signaling. **A)** Kinome activity was quantified of MCF-7 cells treated with 50ng/mL of either FGF2, FGF3, FGF4, FGF10, or FGF19, together with 5µg/mL of heparin. Only the activity dynamics of significantly regulated kinases (biological triplicates, ANOVA p < 0.05) from the PLCγ pathway are plotted. Grey lines represent a 1.5 fold-change, and 90% confidence intervals are presented per quantified phosphorylated peptide.

**Supplementary Figure 6:**
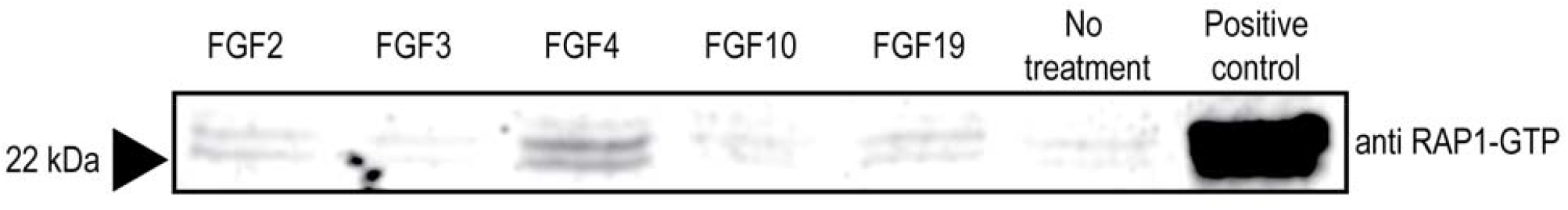
RAP1 pulldown on FGF-stimulated cells. MCF-7 cells were stimulated with either FGF2, FGF3, FGF4, FGF10, and FGF19. Also, a no-stimulation control and a positive control were included. The negative control constituted unstimulated MCF-7 cells. The positive control constituted MCF-7 cell lysate with activated RAP1 by incubating the lysate with GTPγS, which activates all RAP1 in the lysate. The assay consisted of a pulldown of GTP-bound (active) Rap1 of equal amounts of proteins, followed by a western blot using an α–RAP1-GTP antibody.

